# Compatibility and comparative analysis of chronological and biological aging between the legacy 450K and the EPIC v2.0 arrays

**DOI:** 10.1101/2024.10.17.618672

**Authors:** Sven De Pourcq, Pei-Lun Kuo, Ann Zenobia Moore, Stefania Bandinelli, Steve Horvath, Luigi Ferrucci, Valeria Santini

## Abstract

Several different epigenetic clocks built from DNA methylation beadchip arrays, including the Illumina Infinium Human Methylation 450K and EPICv1 BeadChips, have been developed and are used to evaluate biological aging. However, there is still limited information on effectively utilizing the novel EPICv2 arrays and addressing biases in epigenetic clock calculations resulting from missing probes initially present on legacy arrays. We address how the absence of probes on the EPICv2 array, originally present on the 450K array, affects many available epigenetic clock estimates. Using data from the InCHIANTI study of aging, our findings show that while clocks like Hannum and GrimAgeV2 exhibit significant over- or underestimation due to missing probes, the correlations between estimates from all 450K probes (unreduced) and those from shared probes between 450K and EPICv2 (reduced) remain strong. In this cohort, we consider AgeAcceleration from traditional epigenetic clocks to be the most reliable approach. Their values do not significantly change upon probe reduction, with unreduced and reduced values being nearly identical and with stable mean differences across chronological age.

## Short Communication

DNA methylation is a key mechanism regulating chromatin structure and gene expression during development, aging, and cancer. A weighted average of subsets of methylation DNA sites has been used to calculate epigenetic clocks, which estimate biological changes in cells or tissues that occur over time (biological age). The discrepancy between the estimated age by these clocks (DNAmAge) and chronological age provides valuable insights into an organism’s overall health status and aging trajectory (Horvath & Raj, 2018). Most epigenetic clocks are trained using data from DNA methylation beadchip arrays; however, newer versions of these arrays often exclude some of the probes that were available in earlier versions (Noguera-Castells, Garcia-Prieto, Alvarez-Errico, & Esteller, 2023; Peters et al., 2024).

We aim to explore how the absence of certain probes on the EPICv2 array, originally present on older arrays like the 450K, affects the performance of epigenetic clock values. We want to ensure that meaningful biological inferences can still be made across different array versions. Enabling researchers to combine data from different arrays within the same study is becoming increasingly important as the Illumina 450K array is no longer available. Therefore, the EPICv2 array will be the primary choice for future research based on measures of DNA methylation.

We compare four strategies to address this question: (1) using clocks based on all available probes; (2) restricting clocks to probes common to both the 450K and EPICv2 arrays (Dhingra et al., 2019); (3) applying Principal Component (PC) Clock calculations using the full set of probes (Higgins-Chen et al., 2022); and (4) applying PC Clock calculations on probes common to both the 450K and EPICv2 arrays. Comparing these approaches allows us to assess the efficiency and reliability of epigenetic clocks when using the reduced probe set available on the EPICv2 array. A previous study that estimated the Hannum clock between the 450K and EPICv1 arrays provides an example of how applying clock algorithms to a reduced set of the originally included probes can significantly over- or underestimate DNAmAge (Dhingra et al., 2019).

By reducing datasets to account for missing probes, the online DNA Methylation Age Calculator automatically imputes these values for each clock by using the average methylation level of each specific probe from the training dataset. Since the missing probes are the same across individual samples, the imputation process will also be consistent, but the information conveyed by those probes is lost. This method allows us to assess the impact of missing probes on clock estimations directly. The imputation used in PC Clocks is detailed in Supplementary Information.

Here, epigenetic age estimates were first calculated using unreduced 450K data from the InCHIANTI population-based study of aging (n = 732), then recalculated (reduced) using only the shared probes between the EPICv2 and 450K arrays (Table S1). Data preprocessing is detailed in the Supplementary Information. The following epigenetic clocks and their PC counterparts (if available) were evaluated: Horvath1 (Horvath, 2013), Hannum (Hannum et al., 2013), DNAmPhenoAge (Levine et al., 2018), GrimAgeV1 (Lu et al., 2019), GrimAgeV2 (Lu et al., 2022) and DunedinPACE (Belsky et al., 2022). We utilized a modified version of the Illumina manifest to address discrepancies in probe annotation, probe names, and sequences between the EPICv2 manifest and earlier versions (Peters et al., 2024). Out of the 2070 unique probes used to calculate the Horvath1, Hannum, DNAmPhenoAge, GrimAgeV1, GrimAgeV2, and DunedinPACE clocks, only 1821 are included on the EPICv2 (Table 1; details in Supplementary Information).

**Table 1.**
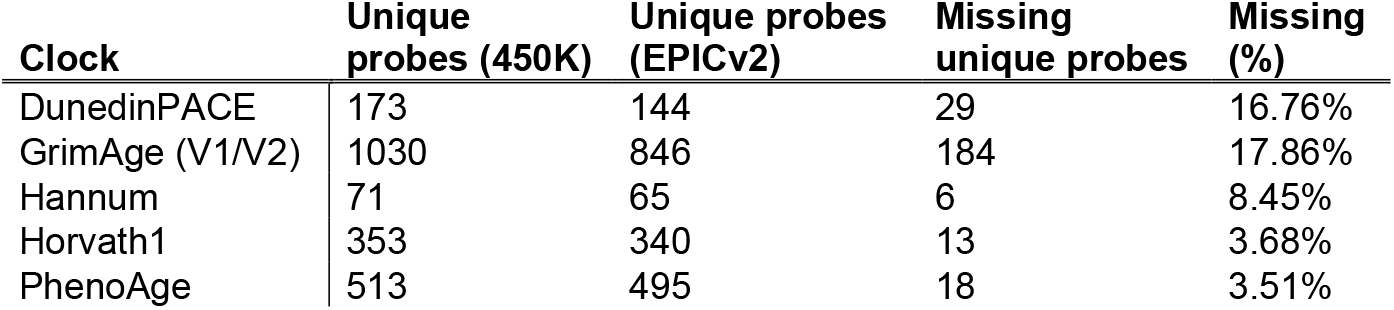
Missing probes on the EPICv2 array per assessed epigenetic clock.

We first compared epigenetic clock estimates from the unreduced 450K InCHIANTI dataset to the reduced dataset, restricted to overlapping probes. Figure 1 illustrates the linear regression and Bland–Altman plots for Horvath1, Hannum, and GrimAgeV2 clocks. Results for the remaining clocks are provided in Supplementary Table S2.

**Figure 1.**
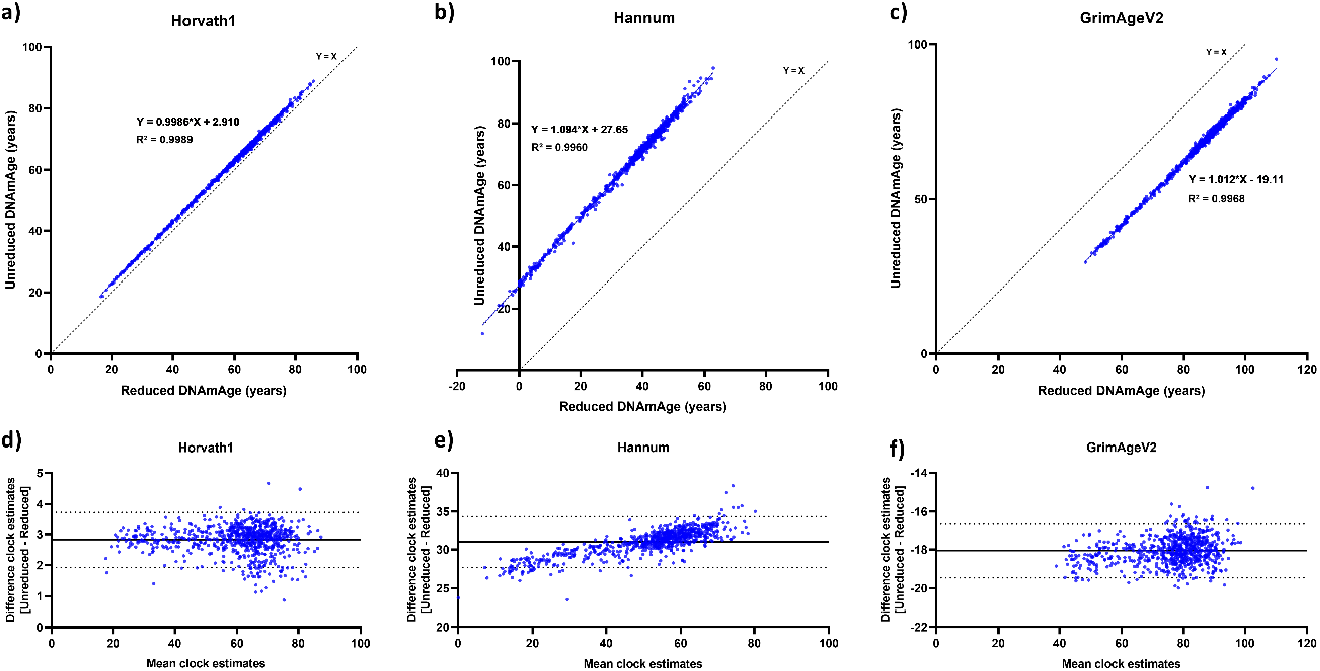
Comparison of epigenetic clock estimates using unreduced vs. reduced 450k array data in the InCHIANTI study of aging. (a) Linear regression with identity line (dotted line) of Unreduced DNAmAge over Reduced DNAmAge for Horvath1; (b) Hannum; and (c) GrimAgeV2. (d) Bland-Altman plots illustrating the differences between the unreduced and reduced measurements against their mean values for Horvath1; (e) Hannum; and (f) GrimAgeV2. The x-axis represents the mean clock estimates of reduced and unreduced datasets combined, while the y-axis shows the difference (unreduced – reduced) between clock estimates. The bold line indicates the mean bias, and the dotted lines represent the ±1.96 standard deviations of the bias. Positive bias values indicate an underestimation for the EPICv2 clock estimates, while negative values indicate an overestimation in clock estimates using the EPICv2.

For the Horvath1 clock, the regression equation was DNAmAge_Unreduced = 0.9986 * DNAmAge_Reduced + 2.91, with a high correlation (Pearson r = 0.9995, p < 0.0001). The mean difference between unreduced and reduced data was 2.83 years (95% CI: 1.38 to 4.27, p < 0.001) (Figure S1a, Table S2). In contrast, the Hannum clock demonstrated significant underestimation in the reduced dataset, with a regression equation of DNAmAge_Unreduced = 1.094 * DNAmAge_Reduced + 27.65 (r = 0.9980, p < 0.0001). The mean difference was much larger at 31.01 years (95% CI: 29.45 to 32.58, p < 0.001), reflecting substantial bias introduced by the missing probes. GrimAgeV2 showed the opposite trend, exhibiting significant overestimation in the reduced dataset. The regression equation was DNAmAge_Unreduced = 1.012 * DNAmAge_Reduced - 19.11 (r = 0.9984, p < 0.0001), with a mean difference of -18.06 years (95% CI: -19.32 to -16.79, p < 0.001). Therefore, missing probes caused consistent overestimation in the reduced dataset for this clock. AgeAcceleration estimates did not show significant differences between the reduced and unreduced data (Figure S1b, Table S2). The Bland–Altman plots showed a bias of 2.83 years (95% limits of agreement (LOA): 1.92 to 3.73) for Horvath1, 31.01 years (95% LOA: 27.68 to 34.35) for Hannum, and -18.06 years (95% LOA: -19.46 to -16.65) for GrimAgeV2 (Figure 1, Figure S4).

Next, we calculated the PC clocks using all available probes. These calculations, designed to reduce technical variation, showed smaller DNAmAge biases than traditional clocks (Figure S1c, Table S2). PCHorvath1 had a minimal bias of 0.1419 years (95% LOA: -1.312 to 1.596). PCHannum exhibited a bias of 0.1060 years (95% LOA: -1.342 to 1.554), suggesting that PC clocks are less sensitive to probe loss.

We then assessed whether the differences between the unreduced and reduced datasets varied with chronological age (Figure S3). For the Hannum clock, we observed a significant relationship (R^2^ = 0.52), indicating that underestimation in the reduced dataset increased with age. For Horvath1 and GrimAgeV2, no significant relationship with age was observed, suggesting stability across age groups. Interestingly, the PC clocks tended to underestimate with increasing chronological age, a phenomenon not seen with the traditional Horvath1 clock but observed in PC Clocks, primarily PCHorvath1, PCHannum, PCPhenoAge, and PCGrimAge (Figure S3c).

Despite the biases observed in traditional clocks, the strong correlations between reduced and unreduced datasets indicate that the biological relevance of epigenetic clocks is largely preserved, even with missing probes. This suggests that reduced datasets, such as those obtained from the EPICv2 array, may still provide reliable associations with clinical outcomes, mortality, and other biological variables.

PC clocks demonstrated minimal DNAmAge bias and potentially offer a robust solution for mitigating the effects of missing probes. They effectively capture broader DNA methylation patterns, making them highly valuable for studies where some probes are unavailable. However, for clocks like GrimAgeV2 that lack PC versions, reliance on reduced datasets may be the only option. Although PC clocks are generally resilient to missing probes, their accuracy may decline with increasing chronological age, as seen with PCHorvath1. Researchers should carefully evaluate the use of PC clocks in studies involving populations with broad age ranges. PC Clocks also remain susceptible to batch effects and may have reduced prediction accuracy compared to original clock outputs (Vavourakis, Herzog, & Widschwendter, 2024). While another study created a new epigenetic clock model to address the issue of missing probes (Garma & Quintela-Fandino, 2024), we opted for using AgeAcceleration from traditional epigenetic clocks. We recommend this approach because reduced AgeAcceleration measures maintain strong correlations with full 450K data and are less variable across chronological age.

Overall, our findings demonstrate that assessing and comparing epigenetic clock values from 450K and EPICv2 arrays is possible with careful data processing.

## Supporting information

Epigenetic Clock Probes EPICv2

## Author Contributions

SDP and VS conceived the study. SDP designed the study, performed data analysis and wrote the draft manuscript. PK contributed to statistical analysis and interpretation. AZM and SH contributed to data interpretation. All authors provided feedback, reviewed, and approved the final manuscript for submission.

## Funding Information

This project has received funding from the European Union’s Horizon 2020 research and innovation programme under the Marie Skłodowska-Curie grant agreement No 953407, and by the Associazione Italiana per la Ricerca sul Cancro (AIRC) IG-26537-2021 Investigator Research Grant. Supported in part by the Intramural Research Program of the National Institute on Aging, NIH, Baltimore, MD, USA.

## Conflict of Interest statement

SH and his team developed the first epigenetic clocks for human saliva, for all tissues (pan-tissue clock), for human mortality risk prediction (PhenoAge, GrimAge), and pan-mammalian clocks. The Regents of the University of California are the sole owner of patents and patent applications directed at epigenetic biomarkers for which SH is a named inventor; SH is a founder and paid consultant of the non-profit Epigenetic Clock Development Foundation that licenses these patents. SH is a Principal Investigator at the Altos Labs, Cambridge Institute of Science, a biomedical company that works on rejuvenation. The remaining authors declare no conflict of interest.

## Data availability statement

Data regarding the InCHIANTI study of aging are available through the website https://www.nia.nih.gov/inchianti-study.

## Supplementary Information

### Data preprocessing

Raw IDATs were processed using the “SeSaMe” package in R (version 4.3.2, sesame 1.20.0). No probes were masked to avoid potential bias from probe imputation during clock calculations. Beta-values were processed using background subtraction with the normal-exponential out-of-band (noob) method. To address inconsistencies in probe annotation, names, and sequences between the EPICv2 and 450K arrays, we used the modified Illumina manifest provided by Peters et al. (2024), specifically the K450locmatch (450K probe names with matching genomic locations in both arrays). This allowed the inclusion of six additional probes for epigenetic clock estimate calculations: cg19693031 (GrimAgeV1/V2), cg01518025 (GrimAgeV1/V2), ch.2.30415474F (Hannum), ch.13.39564907R (Hannum), cg22432269 (Horvath1), and cg05441133 (DNAmPhenoAge). Individual probes needed for each clock were sourced from https://bio-learn.github.io/clocks.html. Clock estimates were calculated using https://dnamage.clockfoundation.org/calculator, and PC Clocks were computed following Higgins-Chen et al. (2022). All analyses were conducted in R version 4.3.2.

### PC Clock imputation

PC clocks utilize principal component analysis and elastic net regression to aggregate data from multiple probes to extract meaningful age-related signals while filtering out noise. This allows PC clocks to capture the essential information conveyed by the clock even if some of the original data are missing. When faced with missing probes, PC clocks handle this by employing mean imputation during the training phase, where each missing value is replaced with the average methylation level of that specific CpG across all samples in the training data.

**Figure S1.**
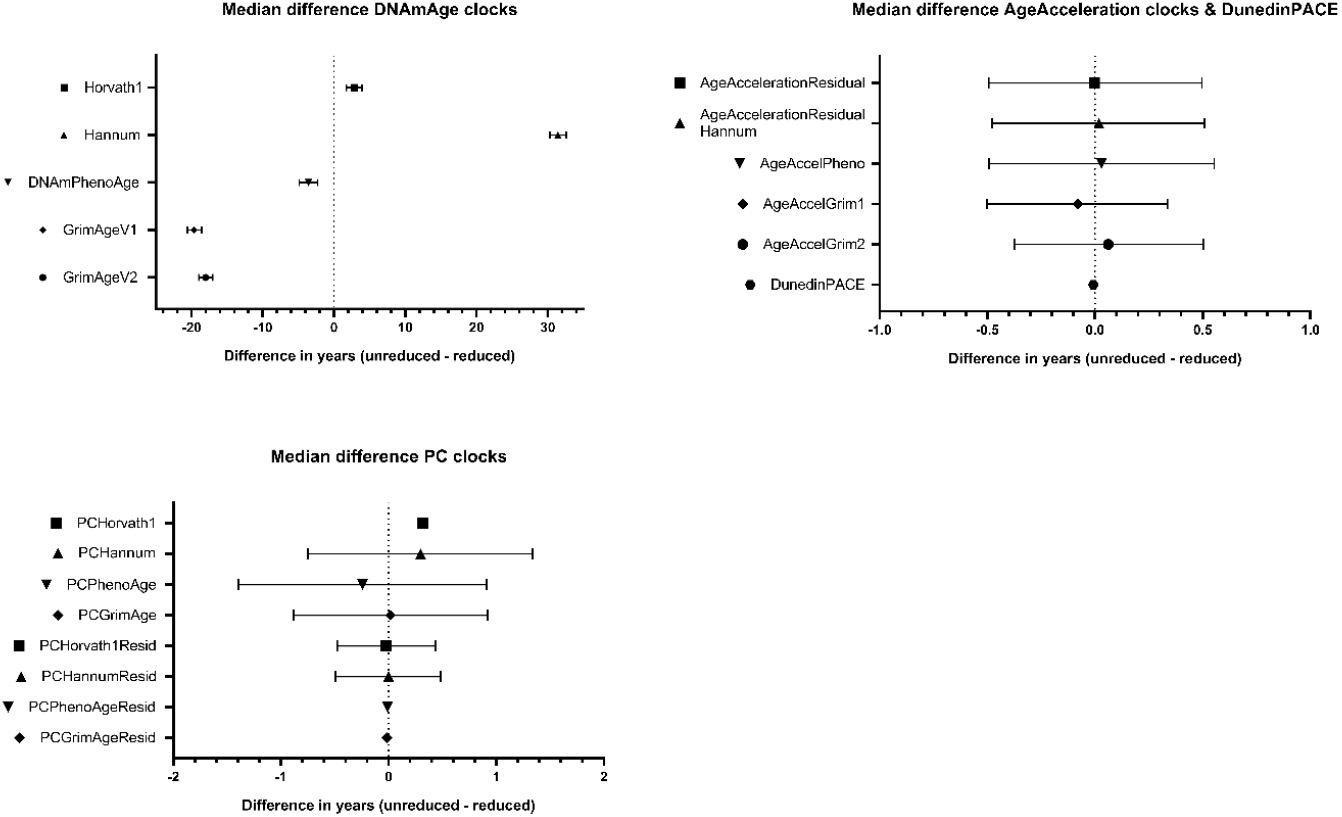
Median differences (unreduced – reduced) in clock estimates for DNAmAge, AgeAcceleration, DunedinPACE and PC Clocks. The median differences between the unreduced and reduced clock estimates. The x-axis depicts the difference in years between clock estimates of the unreduced and reduced data, while the y-axis represents various epigenetic clocks. Whiskers denote the 95% confidence interval. Statistical analysis to assess significant differences between clock estimates from the unreduced and reduced datasets was conducted using Mann-Whitney U analysis.

**Figure S2.**
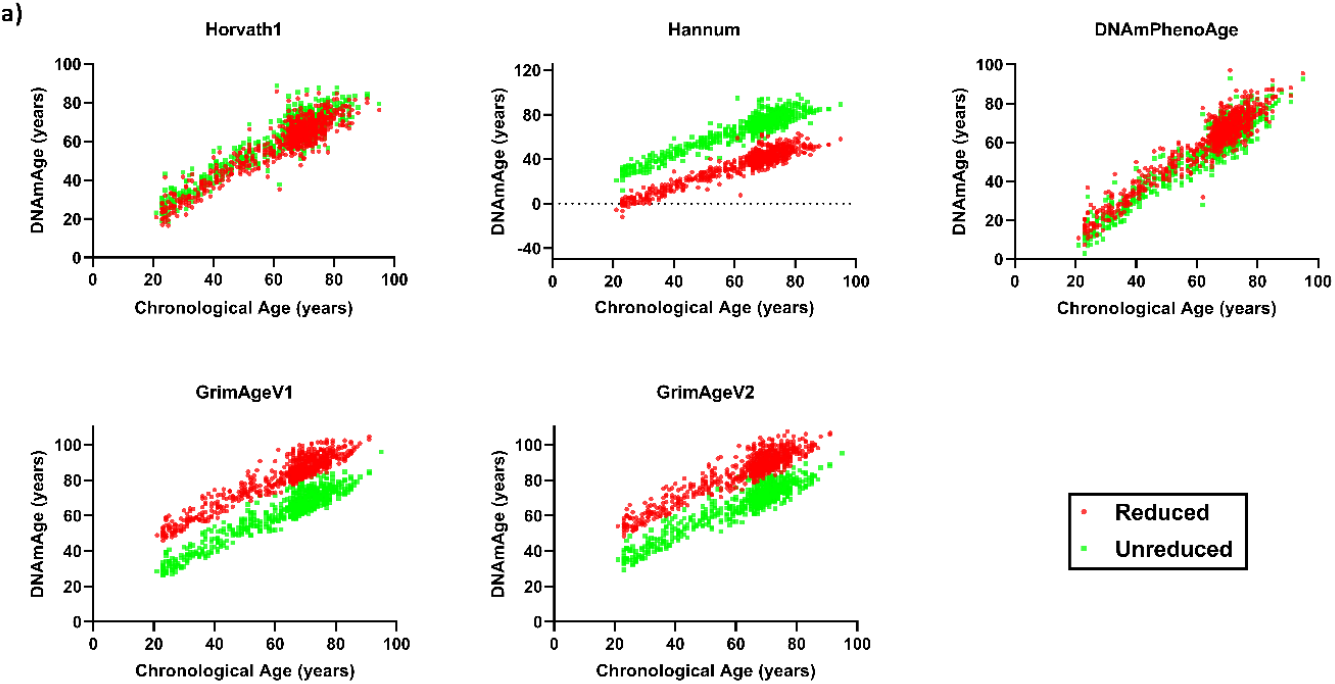
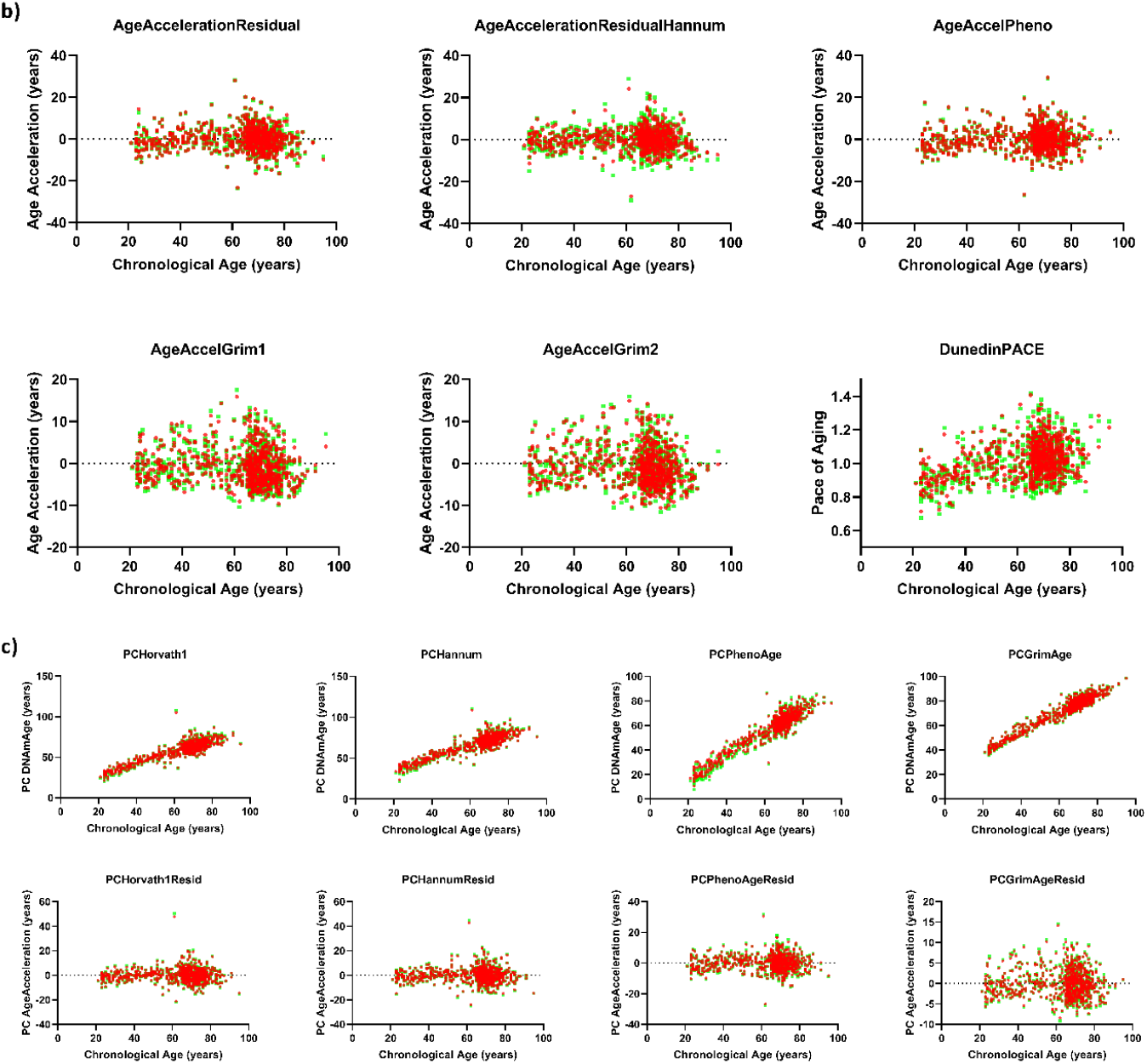
Linear regression of DNAmAge, AgeAcceleration, DunedinPACE and PC Clocks over chronological age. Clock estimates are regressed on chronological age using both reduced and unreduced 450K array data to evaluate whether there is over- or underestimation when increasing chronological age. (a) DNAmAge regressed over chronological age. (b) AgeAcceleration and DunedinPACE regressed over chronological age. (c) PC clocks regressed over chronological age.

**Figure S3.**
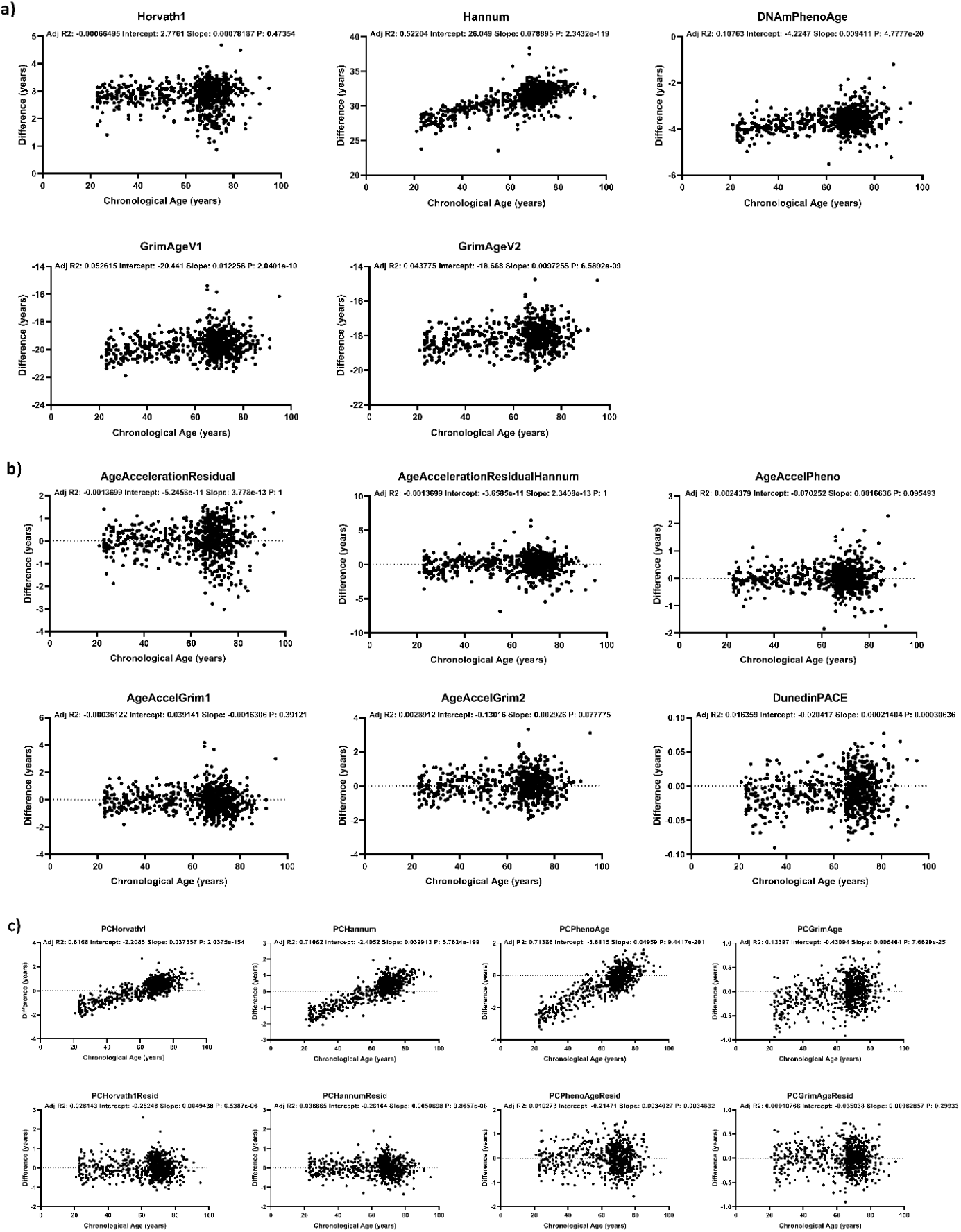
Linear regression of the difference in clock estimates, for DNAmAge, AgeAcceleration, DunedinPACE and PC clocks, over chronological age. Differences in clock estimates (unreduced – reduced) are regressed over chronological age to evaluate whether there is a consistent over- or underestimation across increasing chronological ages. (a) Differences in DNAmAge regressed over chronological age. (b) Differences in AgeAcceleration and DunedinPACE regressed over chronological age. (c) Differences in PC clocks regressed over chronological age.

**Figure S4.**
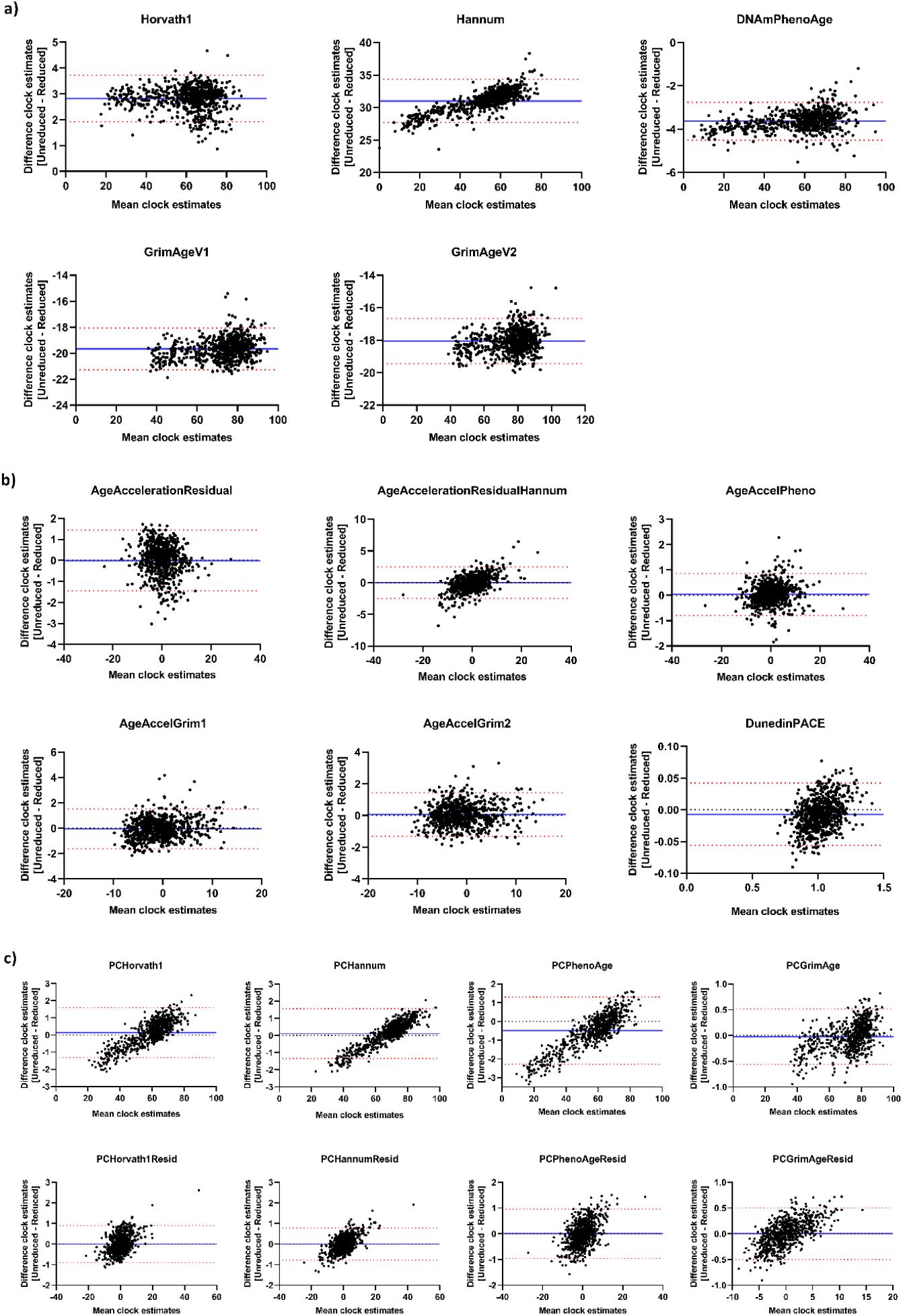
Bland-Altman plots of of DNAmAge, AgeAcceleration, DunedinPACE and PC Clocks. The x-axis represents the mean clock estimates between the two groups, while the y-axis shows the difference (unreduced – reduced) between clock estimates. The blue line indicates the mean bias, and the dotted red lines represent the ±1.96 standard deviations of the bias. Positive bias values indicate an underestimation for the EPICv2 clock estimates, while negative values indicate an overestimation in clock estimates using the EPICv2. (a) Bland-Altman plots of DNAmAge. (b) Bland-Altman plots of AgeAcceleration and DunedinPACE. (c) Bland-Altman plots of PC clocks.

**Table S1.** InCHIANTI study of aging. The age distribution of 732 individuals in the InCHIANTI study, for which DNA methylation data from the 450K array were analysed to calculate epigenetic clock estimates. The table includes mean, standard deviation (SD), median, 25th percentile (p25), 75th percentile (p75), minimum, and maximum ages, provided for the overall cohort as well as separated by sex-at-birth.

|  | n | mean | SD | median | p25 | p75 | min | max |
| --- | --- | --- | --- | --- | --- | --- | --- | --- |
| **Age** | 732 | 62.9 | 15.6 | 68 | 57.8 | 73 | 21 | 95 |
| • **male** | 326 (44.5%) | 61.4 | 15.4 | 67 | 53 | 71 | 23 | 91 |
| • **female** | 406 (55.5%) | 64.1 | 15.7 | 69 | 60.3 | 74 | 21 | 95 |

**Table S2.** Comparison of reduced and unreduced 450K data across epigenetic clocks: differences, linear models, and correlation analysis. A detailed comparison between unreduced 450K and reduced 450K data across various epigenetic clocks. The analysis includes the observed differences between both methods, linear models examining the relationship between reduced (X) and unreduced (Y) datasets, and correlation coefficients assessing the consistency between both approaches. CI: confidence interval; SD: standard deviation; p*: Wilcoxon rank-sum test p-value (two-tailed); p**: Pearson correlation p-value (two-tailed); r: Pearson correlation coefficient.

|  |  |  | Unreduced 450K | Reduced 450K | Difference (Unreduced 450K - Reduced 450K) |  |  | Linear model Reduced 450K (X) vs. Unreduced 450K (Y) |  |  | Correlation |  |
| --- | --- | --- | --- | --- | --- | --- | --- | --- | --- | --- | --- | --- |
|  |  |  | Mean (SD) | Mean (SD) | Mean [95%CI] | Median [95%CI] | p | Slope [95%CI] | Intercept [95%CI] | R<sup>2</sup> | r | p |
| DNAmAge | Clocks | Horvath1 | 61.71 (14.09) | 58.88 (14.11) | 2.825 [1.379, 4.271] | 2.759 [1.726, 3.909] | <0.001 | 0.9986 [0.9962, 1.001] | 2.910 [2.767, 3.053] | 0.9989 | 0.9995 | <0.0001 |
|  |  | Hannum | 66.65 (15.94) | 35.64 (14.54) | 31.01 [29.45, 32.58] | 31.52 [30.29, 32.58] | <0.001 | 1.0940 [1.089, 1.099] | 27.65 [27.46, 27.84] | 0.9960 | 0.9980 | <0.0001 |
|  |  | DNAmPhenoAge | 55.68 (17.36) | 59.31 (17.21) | -3.633 [-5.405, -1.860] | -3.673 [-4.881, -2.289] | <0.001 | 1.009 [1.007, 1.011] | -4.152 [-4.261, -4.043] | 0.9994 | 0.9997 | <0.0001 |
|  |  | GrimAgeV1 | 62.49 (12.87) | 82.16 (12.63) | -19.67 [-20.98, -18.36] | -19.61 [-20.58, -18.62] | <0.001 | 1.017 [1.012, 1.022] | -21.07 [-21.45, -20.69] | 0.9962 | 0.9981 | <0.0001 |
|  |  | GrimAgeV2 | 67.02 (12.43) | 85.08 (12.26) | -18.06 [-19.32, -16.79] | -18.03 [-18.96, -17.05] | <0.001 | 1.012 [1.008, 1.016] | -19.11 [-19.46, -18.75] | 0.9968 | 0.9984 | <0.0001 |
|  | PC Clocks | PCHorvath1 | 58.98 (12.50) | 58.84 (11.88) | 0.1419 [-1.109, 1.392] | 0.2830 [-0.6341, 1.270] | 0.51 | 1.051 [1.049, 1.054] | -2.876 [-3.032, -2.721] | 0.9989 | 0.9994 | <0.0001 |
|  |  | PCHannum | 67.44 (13.19) | 67.34 (12.52) | 0.1060 [-1.213, 1.425] | 0.3846 [-0.7496, 1.340] | 0.58 | 1.053 [1.051, 1.055] | -3.484 [-3.610, -3.359] | 0.9994 | 0.9997 | <0.0001 |
|  |  | PCPhenoAge | 56.73 (15.79) | 57.22 (14.98) | -0.4915 [-2.069, 1.086] | -0.3505 [-1.393, 0.9134] | 0.68 | 1.009 [1.007, 1.011] | -4.152 [-4.261, -4.043] | 0.9994 | 0.9997 | <0.0001 |
|  |  | PCGrimAge | 72.61 (12.62) | 72.63 (12.48) | -0.02425 [-1.311, 1.262] | -0.005960 [-0.8838, 0.9192] | 0.97 | 1.011 [1.010, 1.012] | -0.8261 [-0.9279, -0.7242] | 0.9996 | 0.9998 | <0.0001 |
| AgeAcceleration | Clocks | Horvath1 | -7.377E-08 (5.153) | 2.145E-07 (5.250) | -2.883E-07 [-0.5334, 0.5334] | -0.1246 [-0.4942, 0.4966] | >0.99 | 0.9719 [0.9619, 0.9819] | -2.822E-07 [-0.05238, 0.05238] | 0.9804 | 0.9902 | <0.0001 |
|  |  | Hannum | -5.792E-07 (5.580) | 4.918E-08 (4.883) | -6.284E-07 [-0.5376, 0.5375] | 0.07025 [-0.4787, 0.5087] | 0.94 | 1.119 [1.102, 1.136] | -6.343E-07 [-0.08224, 0.08224] | 0.9588 | 0.9792 | <0.0001 |
|  |  | DNAmPhenoAge | 0.2426 (5.464) | 0.2081 (5.411) | 0.03442 [-0.5231, 0.5920] | 0.03751 [-0.4913, 0.5536] | 0.91 | 1.007 [1.001, 1.013] | 0.03298 [0.002512, 0.06346] | 0.9941 | 0.9971 | <0.0001 |
|  |  | GrimAgeV1 | -0.7528 (4.430) | -0.6894 (4.252) | -0.06345 [-0.5086, 0.3817] | -0.05336 [-0.5027, 0.3363] | 0.70 | 1.025 [1.011, 1.039] | -0.04622 [-0.1047, 0.01223] | 0.9678 | 0.9838 | <0.0001 |
|  |  | GrimAgeV2 | -0.6943 (4.571) | -0.7482 (4.495) | 0.05393 [-0.4109, 0.5187] | 0.09531 [-0.3751, 0.5023] | 0.79 | 1.005 [0.9937, 1.016] | 0.05772 [0.006221, 0.1092] | 0.9766 | 0.9882 | <0.0001 |
|  | PC Clocks | PCHorvath1 | 2.732E-09 (5.371) | -2.650E-07 (5.161) | 2.678E-07 [-0.5401, 0.5401] | -0.08865 [-0.4784, 0.4330] | 0.92 | 1.038 [1.032, 1.044] | 2.777E-07 [-0.03019, 0.03019] | 0.9940 | 0.9970 | <0.0001 |
|  |  | PCHannum | 3.415E-08 (5.829) | -2.828E-07 (5.574) | 3.169E-07 [-0.5848, 0.5848] | -0.09228 [-0.4937, 0.4846] | >0.99 | 1.044 [1.040, 1.048] | 3.295E-07 [-0.02258, 0.02258] | 0.9972 | 0.9986 | <0.0001 |
|  |  | PCPhenoAge | 2.473E-07 (5.103) | -1.366E-08 (4.866) | 2.609E-07 [-0.5112, 0.5112] | -0.1692 [-0.4905, 0.4637] | 0.97 | 1.045 [1.038, 1.051] | 2.615E-07 [-0.03180, 0.03180] | 0.9926 | 0.9963 | <0.0001 |
|  |  | PCGrimAge | -9.563E-09 (3.403) | -1.776E-07 (3.242) | 1.681E-07 [-0.3408, 0.3408] | 0.008325 [-0.3446, 0.3119] | 0.92 | 1.048 [1.043, 1.052] | 1.766E-07 [-0.01470, 0.01470] | 0.9965 | 0.9982 | <0.0001 |
| Pace of Aging | Clocks | DunedinPACE | 1.015 (0.1164) | 1.022 (0.1062) | -0.006951 [-0.01838, 0.004476] | -0.005125 [-0.01912, 0.003921] | 0.20 | 1.073 [1.057, 1.089] | -0.08159 [-0.09836, -0.06481] | 0.9580 | 0.9788 | <0.0001 |

## Notes

https://www.nia.nih.gov/inchianti-study

